# MTS1338, a small *Mycobacterium tuberculosis* RNA, regulates transcriptional shifts consistent with bacterial adaptation for entering into dormancy and survival within host macrophages

**DOI:** 10.1101/474304

**Authors:** Elena G. Salina, Artem Grigorov, Yulia Skvortsova, Konstantin Majorov, Oksana Bychenko, Albina Ostrik, Nadezhda Logunova, Dmitriy Ignatov, Arseny Kaprelyants, Alexander Apt, Tatyana Azhikina

## Abstract

Small non-coding RNAs play a significant role in bacterial adaptation to changing environmental conditions. We investigated the dynamics of expression of MTS1338, a small non-coding RNA of *Mycobacterium tuberculosis*, in the mouse model *in vivo*, regulation of its expression in the *ex vivo* infected macrophages, and the consequences of its overexpression in bacterial cultures. Here we demonstrate that MTS1338 significantly contributes to host-pathogen interactions. Activation of the host immune system triggered NO-inducible up-regulation of MTS1338 in macrophage-engulfed mycobacteria. Constitutive overexpression of MTS1338 in cultured mycobacteria improved their survival *in vitro* under low pH conditions. MTS1338 up-regulation launched a spectrum of shifts in the transcriptome profile similar to those reported for *M. tuberculosis* adaptation to hostile intra-macrophage environment. Using the RNA-seq approach, we demonstrate that gene expression changes accompanying MTS1338 overexpression indicate reduction in translational activity and bacterial growth, which is consistent with entering the dormant state. Taken together, our results suggest a direct involvement on this sRNA in the interplay between mycobacteria and the host immune system during infectious process.

## 1. Introduction

*M. tuberculosis* persistence in the infected host involves several stages and may have different manifestations: initial infection followed by semi-acute or chronic diseases; latent infection characterized by the presence of viable bacteria with slow-to-no level of replication and the lack of clinical manifestations; and transition from the latent state to reactivation processes (Russell, 2007; Stewart et al., 2003). The spectrum of the disease manifestations depends upon a dynamic balance between protective host responses and defensive strategies of *M. tuberculosis*. Identification of molecular mechanisms of *M. tuberculosis* adaptation to the host immune defense during its persistence within macrophages is an important scientific and medical problem.

Long co-evolution of *M. tuberculosis* and its human host allowed the pathogen to develop strategies that can effectively combat host defense systems. Regulatory proteins, non-coding RNAs and their targets constitute complex adaptive metabolic networks that allow the pathogen to resist host response at different stages of infection. Bacterial sRNAs participate in regulation of transcription and translation by affecting the level of gene expression and mRNA stability. Mostly, sRNAs are expressed in response to the external factors, helping bacteria to adaptively react to the changing environmental conditions and regulate the key stages of pathogenesis (Dutta and Srivastava, 2018; Holmqvist and Wagner, 2017; Hor et al., 2018).

Application of the high throughput sequencing and computer algorithm approaches allowed identification of dozens of sRNAs in mycobacterial species (Haning et al., 2014; Schwenk and Arnvig, 2018; Taneja and Dutta, 2019). Several *in vitro* studies have elucidated the functioning of sRNAs in *M. tuberculosis* (Arnvig et al., 2011; Gerrick et al., 2018; Moores et al., 2017; Solans et al., 2014; Mai et al., 2019). However, dissecting the role of a particular sRNA in mycobacterial physiology appeared to be difficult, especially in *in vivo* settings.

One of such RNAs, MTS1338 (DosR-associated sRNA, ncRv11733), is highly expressed during the stationary phase of growth (Arnvig and Young, 2012), and the dormancy state (Ignatov et al., 2015). This sRNA is present only in genomes of highly pathogenic mycobacteria and is very conservative. *In vitro* experiments demonstrated that its transcription is controlled by the transcriptional regulator DosR and is activated under hypoxic and NO-induced stresses (Moores et al., 2017), suggesting that MTS1338 may play a role during the stable phase of infection, when host responses confront mycobacterial multiplication more or less successfully. Indeed, we and others demonstrated a striking increase in the MTS1338 transcription in animal models of chronic infection (Arnvig and Young, 2012; Ignatov et al., 2014). Thus, it seems likely that MTS1338 triggers adaptive biochemical cascades for intracellular persistence.

Here, we characterize the dynamic changes in the MTS1338 expression in mycobacteria obtained from the lungs of genetically susceptible and resistant TB-infected mice, and provide a direct evidence that the level of expression is regulated by the IFN-γ-dependent NO production. Using high-throughput technologies, we describe changes in the genome transcription profile that accompany an increased MTS1338 transcription. Overexpression of MTS1338 leads to transcriptional shifts consistent with decreased bacterial metabolism, cell division and adaptation to host immune responses experienced by mycobacteria residing within host macrophages. Taken together, our results demonstrate that the small non-coding MTS1338 RNA regulates molecular mechanisms providing *M. tuberculosis* inter-macrophage survival.

## 2. Materials and Methods

### Bacterial strains, media and growth conditions

For *in vitro* experiments, *M. tuberculosis* H37Rv, pMV (empty plasmid control) and OVER (MTS1338 overexpressing) *M. tuberculosis* strains were initially grown from frozen stocks for 10 days in Sauton medium. Medium content (per liter): 0.5 g KH_2_PO_4_, 1.4 g MgSO_4_×7H_2_O, 4 g L-asparagine, 60 ml glycerol, 0.05 g ferric ammonium citrate, 2 g sodium citrate, 0.1 ml 1% ZnSO_4_, pH 7.0 (adjusted with 1M NaOH). Supplements: ADC growth supplement (Connell, 1994), 0.05% Tween 80 and 50 µg/ml kanamycin (Sigma-Aldrich, USA). Growth conditions: 37°C with agitation (200 rpm). The starter cultures were inoculated into fresh medium (the same composition) and grown up to stationary phase for RNA-seq experiments and stress survival experiments. For cloning procedures, *Escherichia coli* DH5α was grown in Luria Bertani (LB) broth and LB-agar. When required, antibiotics were added at the following concentrations: kanamycin (Sigma-Aldrich), 50 μg/ml (*M. tuberculosis*); ampicillin (Invitrogen, USA), 100 μg/ml (*E. coli*).

### M. tuberculosis OVER and pMV (control) strains establishment

The MTS1338 gene-containing vector was constructed on the basis of the pMV261 (Stover et al., 1991) as described by (Ignatov et al., 2015). The plasmid was transferred into mycobacteria by electroporation. MTS1338 overexpression was confirmed by quantitative PCR. The control strain was produced using an empty pMV261 vector.

### Growth inhibition *in vitro* by NO, H_2_O_2_ and low pH

Bacterial cultures were grown up to the stationary phase, washed up with PBS and diluted to OD_600_=0.2 (10^7^ CFU/ml) by (i) Sauton medium (pH 5.5) with ADC growth supplement and 0.05% Tween for low pH stress; (ii) by the culture supernatant to study inhibitory effects of NO (provided by the DETA/NO donor, 0.5 mM) and H_2_O_2_ (10 mM). Cell viability after 24 h and 48 h of stresses exposure were measured by incorporation of [^3^H]-uracil label. 2 μl 5,6,-[^3^H]-uracil (2 μCi) were added to 1-ml culture samples and incubated at 37°C with agitation for 20 h. 200μl of culture were put in 3 ml 7% ice-cold CCl_3_COOH, incubated at 0° C for 15 min and filtered through glass microfiber filters (Whatman, USA). Precipitated cells were washed with 3 ml 7% CCl_3_COOH and 3 ml 96% ethanol. Filters were put in 10 ml of scintillation mixture; CPM were determined by LS analyser (Beckman Instruments Inc, USA).

### RNA extraction from cultured mycobacteria

Bacterial cultures were grown up to the stationary phase, rapidly cooled on ice, centrifuged, and total RNA was isolated by phenol-chloroform extraction after cell disruption with Bead Beater (BioSpec Products, USA) as previously described (Rustad et al., 2009). After isolation, RNA was treated with Turbo DNase (Life Technologies, USA) to remove traces of genomic DNA, and purified with the RNeasy mini kit (Qiagen, Netherlands). Amounts and purity of RNA were determined spectrophotometrically; integrity of RNA was assessed in 1% agarose gel.

### Libraries for RNA-seq and RNA-seq data analyses

RNA samples were depleted of 16S and 23S rRNA using RiboMinus™ Transcriptome Isolation Kit, bacteria (Invitrogen, USA). Sequencing libraries were generated using the resulting ribosomal transcript-depleted RNA and NEBNext Ultra II Directional RNA Library Prep Kit (NEB, USA) according to the manufacturers’ protocol. Sequencing was performed using the Illumina NovaSeq as the single-ended 100 nt-long reads. Experiments were performed in triplicates.

After quality control evaluation and trimming of bad qualitative reads the reads were mapped on the reference *M. tuberculosis* genome (AL123456.3, http://www.ncbi.nlm.nih.gov/) by Bowtie2 (Langmead and Salzberg, 2012). The alignment was performed with the “-local” option, which allows leaving 5’ and 3’ ends uncharted. Calculation of the mapped reads for all genes was performed using functions of the featureCounts package (Liao et al., 2014) built into the author’s script. Resulting statistics were visualized as transcription profiles using the Artemis genome browser (Carver et al., 2012).

Differentially expressed genes were identified by the software package DESeq2 (Love et al., 2014). The genes were considered to be differentially expressed, if the p-value was less than 0.05, the expected measure of false deviations (FDR) was not higher than 0.1, and the expression change module (FC, Fold change) was not less than 3. Further distribution of genes according functional categories was performed using the Mycobrowser database (https://mycobrowser.epfl.ch/).

### Quantitative reverse transcription-PCR (qRT-PCR)

One microgram of total RNA was used for cDNA synthesis with random hexanucleotides and SuperScript III reverse transcriptase (Life Technologies, USA). Quantitative PCR was performed using qPCRmix-HS SYBR (Evrogen, Russia) and the Light Cycler 480 real-time PCR system (Roche, Switzerland); cycling conditions were as follows: 95ºC for 20 s, 61ºC for 20 s, 72ºC for 30 s, repeat 40 times; primers are listed in Suppl Table 1. In the end of amplification, a dissociation curve was plotted to confirm specificity of the product. All real-time experiments were repeated in triplicate. The results were normalized against the 16S rRNA gene. Calculations were performed according to (Ganger et al., 2017) for the relative expression ratio.

**Table 1.** List of genes, differentially expressed in OVER strain *vs* pMV strain

| *Gene* |  | Function (according to Mycobrowser) | Functional category | Essentiality in vitro (Griffin et al., 2011; Sasseti et al., 2003) | Fold Change |
| --- | --- | --- | --- | --- | --- |
| *Rv0079* |  |  | Conserved hypotheticals |  | 8,2 |
| *Rv0080* |  |  | Conserved hypotheticals |  | 10,0 |
| *Rv0081* |  |  | Regulatory proteins |  | 4,9 |
| *Rv0082* |  | Probable oxidoreductase | Intermediary metabolism and respiration |  | 6,1 |
| *Rv0083* |  | Probable oxidoreductase | Intermediary metabolism and respiration |  | 5,7 |
| *Rv0084* | *hycD* | Possible formate hydrogenlyase HycD | Intermediary metabolism and respiration |  | 4,0 |
| *Rv0085* | *hycP* | Possible hydrogenase HycP | Intermediary metabolism and respiration | ES | 6,2 |
| *Rv0086* | *hycQ* | Possible hydrogenase HycQ | Intermediary metabolism and respiration | ES | 5,2 |
| *Rv0087* | *hycE* | Possible formate hydrogenase HycE | Intermediary metabolism and respiration |  | 4,2 |
| *Rv0440* | *groEL2* | 60 kDa chaperonin 2 GroEL2 (protein CPN60-2) (GroEL protein 2) (65 kDa antigen) (heat shock protein 65) (cell wall protein A) (antigen A) | Virulence, detoxification, adaptation | ES | 0,2 |
| *Rv0516c* |  | Possible anti-anti-sigma factor | Information pathways |  | 2,9 |
| *Rv1158c* |  | Conserved hypothetical ala-, pro-rich protein | Conserved hypotheticals |  | 0,05 |
| *Rv1469* | *ctpD* | Probable cation transporter P-type ATPase D CtpD | Cell wall and cell processes |  | 0,4 |
| *Rv1620c* | *cydC* | Probable 'component linked with the assembly of cytochrome' transport transmembrane ATP-binding protein ABC transporter CydC | Intermediary metabolism and respiration | ES | 7,4 |
| *Rv1621c* | *cydD* | Probable 'component linked with the assembly of cytochrome' transport transmembrane ATP-binding protein ABC transporter CydD | Intermediary metabolism and respiration |  | 5,7 |
| *Rv1622c* | *cydB* | Probable integral membrane cytochrome D ubiquinol oxidase (subunit II) CydB (cytochrome BD-I oxidase subunit II) | Intermediary metabolism and respiration | ES | 5,7 |
| *Rv1690* | *lprJ* | Probable lipoprotein LprJ | Cell wall and cell processes |  | 0,3 |
| *Rv1772* |  |  | Conserved hypotheticals |  | 0,3 |
| *Rv2033c* |  |  | Conserved hypotheticals |  | 0,2 |
| *Rv2812* |  | Probable transposase | Insertion seqs and phages |  | 0,1 |
| *Rv2947c* | *pks15* | Probable polyketide synthase Pks15 | Lipid metabolism |  | 0,2 |
| *Rv2986c* | *hupB* | DNA-binding protein HU homolog HupB (histone-like protein) (HLP) (21-kDa laminin-2-binding protein) | Information pathways | ES | 0,2 |
| *Rv2987c* | *leuD* | Probable 3-isopropylmalate dehydratase (small subunit) LeuD (isopropylmalate isomerase) (alpha-IPM isomerase) (IPMI) | Intermediary metabolism and respiration | ES | 0,2 |
| *Rv2988c* | *leuC* | Probable 3-isopropylmalate dehydratase (large subunit) LeuC (isopropylmalate isomerase) (alpha-IPM isomerase) (IPMI) | Intermediary metabolism and respiration |  | 0,3 |
| *Rv3019c* | *esxR* | Secreted ESAT-6 like protein EsxR (TB10.3) (ESAT-6 like protein 9) | Cell wall and cell processes |  | 0,3 |
| *Rv3136* | *PPE51* | PPE family protein PPE51 | Pe/ppe |  | 0,1 |
| *Rv3260c* | *whiB2* | Probable transcriptional regulatory protein WhiB-like WhiB2 | Regulatory proteins | ES | 0,2 |
| *Rv3767c* |  | Possible S-adenosylmethionine-dependent methyltransferase | Lipid metabolism |  | 0,3 |

### Infections *in vivo* and ex vivo

#### Mycobacteria

For infection of mice and macrophage cultures, *M. tuberculosis* H37Rv (substrain Pasteur) from the collection of CIT were used. Mycobacteria were prepared to infect mice and macrophages as described previously (Lyadova et al., 2000). Briefly, to obtain log-phase bacteria for challenge, 50 µl from a thawed aliquot was added to 30 ml of Dubos broth (BD Biosience, USA) supplemented with 0.5% Fatty Acid-Poor BSA (Calbiochem-Behring Corp., USA) and oleic acid and incubated for 2 weeks at 37°C. The resulting suspension was washed two times at 3000 g, 20 min, 4°C with Ca^2+^- and Mg^2+^-free PBS containing 0,2 mM EDTA and 0,025% Tween 80. Cultures were filtered through a 45 µm-pore-size filter (Millipore, USA) to remove clumps. To estimate the CFU content in the filtrate, 20 µl from each 5-fold serial dilution was plated onto Dubos agar (BD), and the total number of micro-colonies in the spot was calculated under an inverted microscope (200^x^ magnification) after being cultured for 3 days at 37°C. The bulk of the filtered culture was stored at 4°C, and it was found that no change in the CFU content occurred during this storage period.

#### Mice

C57BL/6Ycit (B6) and I/StSnEgYCit strain (I/St) mice were kept under conventional, non-SPF conditions in the Animal Facilities of the Central Research Institute of Tuberculosis (CIT, Moscow, Russia) in accordance with the guidelines from the Russian Ministry of Health # 755, and under the NIH Office of Laboratory Animal Welfare (OLAW) Assurance #A5502-11. Female mice aged 2.5–3.0 months were used. All experimental procedures were approved by the Bioethics Committee of the Central Research Institute of Tuberculosis (IACUC), protocols # 2, 3, 7, 8, 11 approved on March 6, 2016.

#### Infection of mice

To infect mice, mycobacteria were re-suspended in supplemented PBS. Mice were infected via respiratory tract with ~100 viable CFU/mouse using an Inhalation Exposure System (Glas-Col, USA), as described in (Radaeva et al., 2008; Radaeva et al., 2005). The size of challenging dose was confirmed in preliminary experiments by plating serial 2-fold dilutions of 2-ml homogenates of the whole lungs obtained from B6 and I/St females at 2 h post-exposure onto Dubos agar and counting colonies after 3-wk incubation at 37ºC. To assess CFU counts, lungs from individual mice were homogenized in 2.0 ml of sterile saline, and 10-fold serial dilutions were plated on Dubos agar and incubated at 37°C for 20-22 days.

#### Infection of peritoneal macrophages, iNOS activation, RNA extraction

To obtain peritoneal macrophages, B6 mice were injected intra-peritoneally with 3% peptone (Sigma-Aldrich) in saline. Five days later, peritoneal exudate cells (PEC) were eluted from the peritoneal cavities with Ca^2+^- and Mg^2+^-free PBS supplemented with 2% FCS and 10 U/ml heparin, washed twice with PBS, and resuspended in RPMI 1640 containing 5% FCS, 10 mM HEPES and 2 mM L-glutamine. The content of nonspecific esterase-positive cells in PEC exceeded 85 %. PEC were plated onto 90 mm Petri dishes (Costar, Corning Inc., USA) at 10 × 10^6^ cells/dish in 10 ml of RPMI-1640 containing 5% FCS, 10 mM HEPES and 2 mM L-glutamine to obtain macrophage monolayers. The cells were allowed to adhere for 2 h at 37°C, 5% CO_2_ before mycobacteria were added in 10 ml of supplemented RPMI-1640 at MOI = 30, 20, 15 and 5 for further culturing for periods indicated in Figure 2. Macrophage-free mycobacterial cultures served as controls.

To activate macrophages, monolayers were treated with murine rIFN-γ (100 U/ml, Sigma) for 14 h before adding mycobacteria. To block iNOS, 100 µM L-NIL (Sigma) was added 1 h before rIFN-γ administration.

To extract RNA, dishes with cell monolayers were gently shaken, culture medium was completely aspirated and macrophages were lysed with 5 ml/dish of Trizol (Invitrogen) as recommended by the manufacturer. Mycobacteria alone in control cultures were suspended by pipetting and centrifuged at 3000 g, 20 min, 4°C. Pellets were suspended in 1 ml of Trizol.

#### Statistics

Statistical analysis was performed using ANOVA test and t-test by GraphPad Prism6.0 software (GraphPad Software, San Diego, CA, USA). *P* < 0.05 was considered statistically significant.

## 3. Results

### 3.1 MTS1338 expression in TB-infected mice

Earlier it was demonstrated that several *M. tuberculosis* non-coding RNAs, including MTS1338, are highly transcribed *in vivo* (Arnvig et al., 2011; Ignatov et al., 2014). Here, we investigated the dynamical transcription profile of MTS1338 in mycobacteria extracted from the mouse lungs from initial to terminal phases of infection. Aerosol infection with low doses of *M. tuberculosis* leads to a chronic and temporary effectively controlled infection in genetically resistant B6 mice, whilst in susceptible I/St mice fatal pulmonary pathology develops relatively rapidly (Kondratieva et al., 2010). Differences in mycobacterial lung CFU counts between I/St and B6 mice reached about 1.2 logs during the first 2 months post challenge and remained stable until I/St mice succumbed to infection (Figure 1A). We profiled the MTS1338 expression in the lung mycobacterial population by quantitative real-time PCR (Figure 1B). The highest level of expression was observed at week 10 post-challenge. In B6 mice, it remained high throughout the experiment, although slowly decreased at the very late phase of infection. At week 10 of infection, when I/St mice start to lose control of the disease progression, the level of MTS1338 expression in their lung mycobacterial population was significantly higher (*P* < 0.01) than that in more resistant B6 mice (Figure 1B). Overall, at the stage of flourishing infection, the MTS1338 expression level in the lung-residing bacteria was more then 1000-fold higher compared to its expression level during stationary phase of growth *in vitro* (Ignatov et al., 2014).

**Figure 1.**
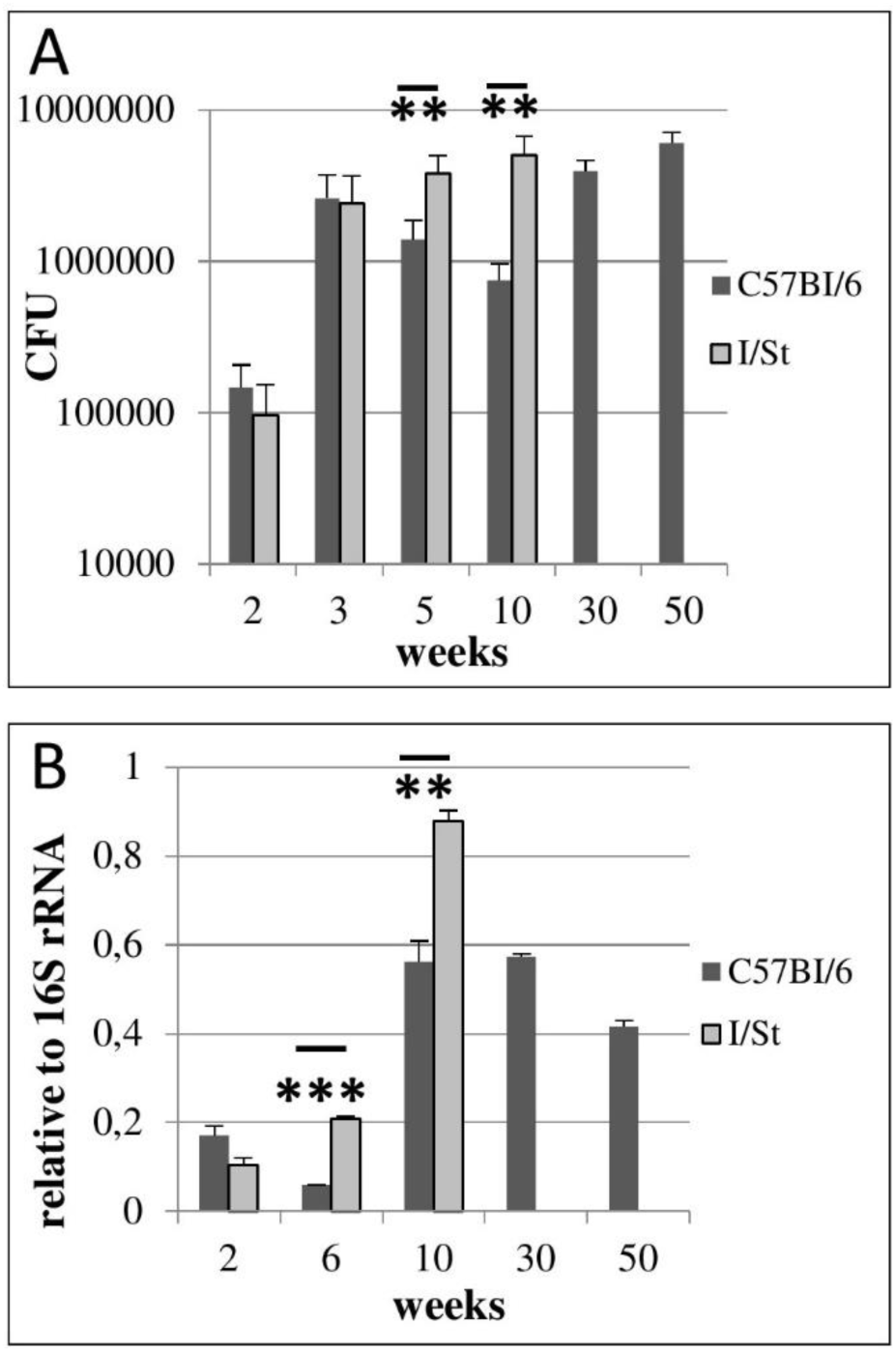
*M. tuberculosis* infection in resistant B6 and susceptible I/St mice. (A). Lung CFU counts along the disease progression (\*\**P* < 0.01 at 5- and 10-weeks post challenge, ANOVA). (B). MTS1338 expression levels of at different time points. (\*\**P* < 0.01 and \*\*\**P* < 0.001, unpaired *t*-test). At indicated time points, samples of total RNA were analyzed by quantitative real-time PCR, and the MTS1338 expression levels in the lung tissue were normalized to those of 16S rRNA. The data are presented as the mean ± SD of three independent experiments.

### 3.2 The expression of MTS1338 is regulated by iNOS

Our *in vivo* experiments demonstrated that the level of MTS1338 expression peaks at the stage of fully developed adaptive immune response against mycobacteria. At this stage, B6 mice display significantly higher levels of IFN-γ production compare to their I/St counterparts (Logunova et al., 2015; Radaeva et al., 2005). Since IFN-γ is the key cytokine activating macrophages for intracellular mycobacterial killing (Cooper, 2009), we compared MTS1338 expression levels in infected peritoneal B6 macrophages, either activated by the external IFN-γ, or not. The level of MTS1338 expression was assessed in dynamics at 2, 4, and 24 hours of macrophage infection (Fig. 2A). In IFN-γ-activated macrophages, MTS1338 expression was significantly (*P* < 0.001, unpaired *t*-test) higher than in control macrophages at every time point, and the difference reached more than 10-fold at 24 hours post infection. Thus, pre-activation of macrophages with IFN-γ induced up-regulation of the MTS1338 expression in engulfed mycobacteria. Given that the efficacy of mycobacterial killing by peritoneal macrophages significantly increases in the presence of IFN-γ (Majorov et al., 2003), this result suggests that the level of MTS1338 expression correlates with the level of pressure emanating from macrophage antibacterial systems.

**Figure 2.**
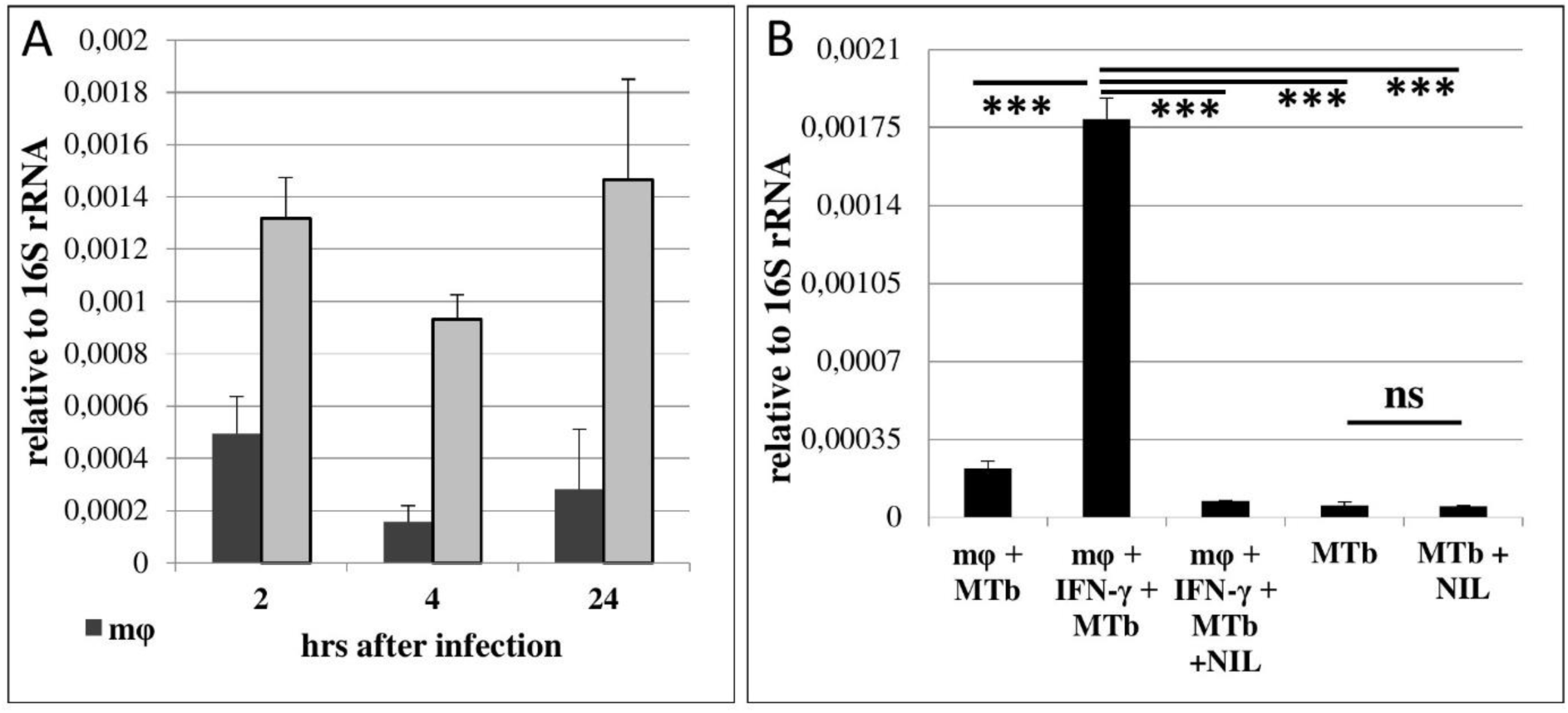
MTS1338 transcription is NO-dependent and correlates with activation of infected macrophages. (A). The MTS1338 transcription dynamics in infected peritoneal macrophages of B6 mice. (B). The level of MTS1338 transcription at 24 h post infection: control (mφ +MTb), IFN-γ-activated (mφ +MTb + INF-γ), IFN-γ-activated and L-NIL treated (mφ +MTb + INF-γ + NIL). The levels of MTS1338 transcription in pure *M. tuberculosis* cultures (MTb) and L-NIL-treated cultures (MTb + NIL) serve as controls for the assessment of possible L-NIL influence onto cultured mycobacteria. The data are presented as the mean ± SD of three independent experiments; \*\**P* < 0.01, \*\*\**P* <0.005, ns – not significant, unpaired *t*-test).

Since the active nitrogen oxidative derivatives serve as the major trigger of MTS1338 transcription activation *in vitro* (Moores et al., 2017), we decided to test whether this is true for the infected macrophage system. Nitrogen oxidative derivatives production in macrophages depends upon inducible NO-synthase (iNOS2), thus we compared mycobacteria-infected IFN-γ-activated and control macrophages cultured for 24 hours in the presence or absence of L-NIL [N6-(1-iminoethyl)-L-lysine hydrochloride] – a selective inhibitor of iNOS2. Inhibition of NO production in IFN-γ-activated macrophages completely abrogated elevation in the MTS1338 expression. L-NIL itself did not affect MTS1338 expression in pure *M. tuberculosis* cultures (Figure 2B). Thus, in macrophages, nitrogen oxidative derivatives are an important trigger of MTS1338 expression.

### 3.3 Survival under *in vitro* stresses

To check whether elevated transcription of MTS1338 protects *M. tuberculosis* against hostile stressful environment, we compared survival of the OVER and control strains in cultures subjected to different type of stresses: low pH or elevated levels of NO and H_2_O_2_. Inhibitory effects of external NO, H_2_O_2_ and pH = 5.5 on mycobacteria were estimated by the level of incorporation of [^3^H]-uracil after 24 and 48 h of stress exposure (Figure 3). In the absence of stress, the OVER strain grew slightly slower than the control one (*P* < 0.01 at the 48-h time point), which is consistent with earlier observations (Arnvig et al., 2011; Ignatov et al., 2015). Treatment of cultures with external NO or H_2_O_2_ had marginal to no effect on mycobacterial growth. However, overexpression of MTS1338 provided significant level of protection against acidic conditions: at pH = 5.5, uracil incorporation by the OVER strain was significantly higher both at 24 h (*P* < 0.05), and 48 h (*P* < 0.01) of culturing.

**Figure 3.**
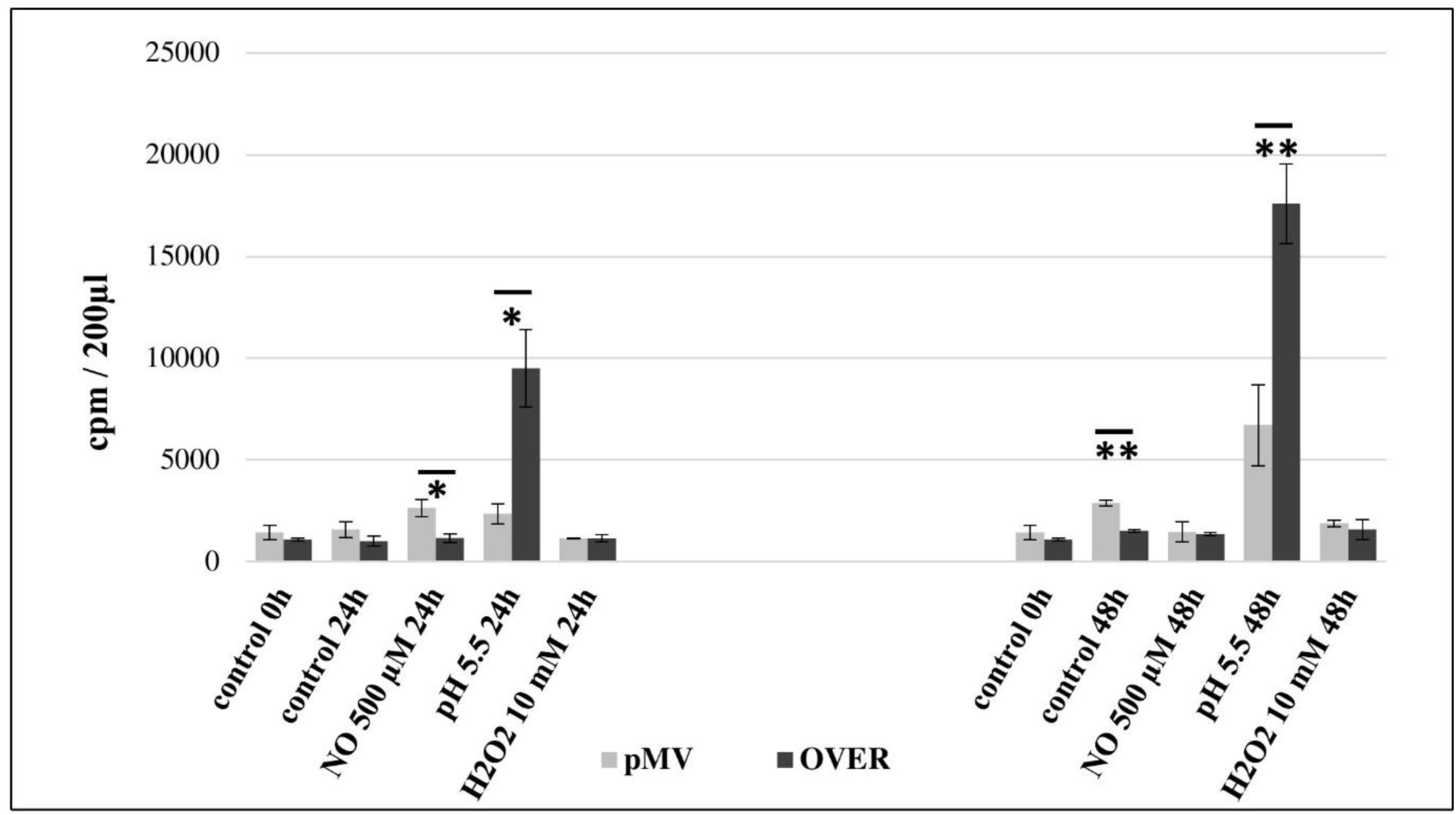
Viability of the OVER and control *M. tuberculosis* strains under stressful conditions in vitro. Stationary phase mycobacteria were subjected to pH = 5.5 or elevated levels of NO and H_2_O_2_ in 24-h and 48-h cultures. The effect of stresses was measured by [^3^H]-uracil incorporation in three independent experiments and expresses as mean CPM ± SD. \**P* < 0.05, \*\**P* < 0. 01, unpaired *t*-test). The data are presented as the mean ± SD of three independent experiments.

### 3.4 Transcriptome changes induced by the MTS1338 overexpression are consistent with mycobacterial adaptation to persistence

To assess how overexpression of MTS1338 influences mycobacterial adaptation, we compared transcriptomes of the OVER and control pMV strains at the phase of stationary growth in liquid culture using RNA-seq approaches. The MTS1338 expression level in the OVER strain in these experiments was more than 10-fold higher compared to the pMV strain as confirmed by qRT-PCR (Suppl Figure 1A).

Mapping the processed reads against the reference *M. tuberculosis* genome (AL123456.3, http://www.ncbi.nlm.nih.gov/), provided the following numbers of mapped reads: 18940746 (96.91%), 19931958 (97.27%) and 20528207 (97.68%) for the OVER strains and 18671706 (97.04%), 16003598 (96.36%) and 15540058 (96.85%) for the pMV strain. The percentage of the protein-encoding part of the genome deduced from all mapped reads comprised 13.97% (2609191), 20.04% (3207752) and 15.52% (2411509) for pMV, and 10.9% (2065273), 15.39% (3067710) and 20.11% (4127293) for OVER (12.8 × 10^6^ reads).

Using the software package DESeq2 (Love et al., 2014), we identified genes the expression of which differed between the two strains. Unexpectedly, only 28 genes were found to change their expression more then 1.5-fold under the MTS1338 overexpression conditions, with 15 genes demonstrating a decreased and 13 genes an increased expression. Further ascribing of genes to functional categories was performed using the Mycobrowser database. The list of differentially expressed genes (DEGs) is displayed in Table 1. Complete data on RNA-seq are displayed in Suppl. Table 2. Possible functional consequences of particular shifts in gene expression profiles are provided in the Discussion section. Differential expression of six randomly chosen genes was confirmed by the quantitative RT-PCR (Suppl. Figure 1B).

## 4. Discussion

In mycobacteria, sRNAs have been discovered much later then in many other bacterial species (Haning et al., 2014), and their functions mostly remain unknown. However, recent high-throughput transcriptional profiling of cultured *M. tuberculosis* exposed to relevant stresses identified a pool of both known and novel mycobacterial sRNAs involved in response to stress conditions *in vitro* (Gerrick et al., 2018).

Here, we present functional characteristics of the sRNA MTS1338, one of highly expressed in *M. tuberculosis* during the stationary growth phase (Arnvig et al., 2011) and at dormancy (Ignatov et al., 2015), suggesting its role in the maintenance of *M. tuberculosis* survival under unfavorable conditions. Since these observations suggest that high levels of MTS1338 expression are required for its functional activity, we constructed the *M. tuberculosis* strain overexpressing MTS1338 for identification of its *in vitro* phenotype, as well as transcriptional changes triggered by this small RNA. Earlier it was demonstrated that MTS1338 expression is NO-inducible and is activated by transcriptional regulator DosR under hypoxic cultural conditions (Moores et al., 2017), as well as under starvation, oxidative and low pH stresses (Gerrick et al., 2018). Our experiments demonstrate that the strain constitutively overexpressing MTS1338 is more resistant to low pH than the control strain (Figure 3).

Overexpression of MTS1338 dramatically changes the bacterial growth rate (Arnvig et al., 2011; Ignatov et al., 2015). Our experiments with *M. tuberculosis* dormancy and resuscitation *in vitro* demonstrated that MTS1338 participates in entering dormancy (Ignatov et al., 2015), but is not involved in the resuscitation process (Salina et al., 2019). *In vivo*, high levels of MTS1338 transcription were reported for *M. tuberculosis* residing in chronically infected mouse lungs (Arnvig et al., 2011; Ignatov et al., 2014). In the present work, using a mouse model of infection, we demonstrate that MTS1338 up-regulation strictly follows activation of iNOS in macrophages. Importantly, at the stage of advanced infection the level of expression was significantly higher in genetically TB-susceptible I/St mice compared to more resistant B6 animals. This may reflect an attempt of mycobacteria residing in the I/St lungs to rapidly turn down metabolism, facing severe functional failure in the surrounding tissue, providing aggressive, highly hypoxic and necrotic conditions to a large proportion of mycobacterial population (Kondratieva et al., 2010). We anticipated that an abundant expression of MTS1338 leads to shifts of the whole genome transcriptional profile towards preparation of mycobacteria to stress-induced metabolic slowdown, thereby helping survival in hostile intra-macrophage surrounding

However, RNA-seq transcriptome evaluation demonstrated that the number of DEGs in MTS1338-overexpressing and control strains is relatively small. MTS1338 overexpression resulted in elevated in the expression of three operons – *Rv0079-Rv0081*, *Rv0082-Rv0087* and *Rv1620c-Rv1622c*. *Rv0079-0081* genes belong to the DosR regulon which activates under hypoxic conditions (Voskuil et al., 2003). *Rv0079* expression was shown to be regulated by *Rv0081* (Chauhan et al., 2011). In *E. coli* and *M. bovis*, homologous protein significantly inhibits cell growth, apparently interacting with the 30S ribosome subunit and inhibiting translation – the phenotype typical for transition to dormancy (Kumar et al., 2012). *Rv0080* encodes a conservative hypothetical protein with unknown functions. It contains a domain of pyridoxine 5’-phosphate (PNP) oxidase-like (PNPOx-like) superfamily, which catalyze flavin mononucleotide-mediated redox reactions. *Rv0081* is one of two key transcriptional factors mediating early response to hypoxia (Galagan et al., 2013). As an important “metabolic hub” working in concert with other transcription regulators, *Rv0081* is associated with the processes of lipid metabolism, protein degradation and cholesterol biosynthesis.

The genes of other two operons encode proteins of the functional category “Intermediary metabolism and respiration”. *Rv0082-Rv0087* genes are also regulated by *Rv0081* (He et al., 2011), but not included in the DosR regulon. The *Rv0082–Rv0087* locus in *M. tuberculosis* encodes a putative [NiFe]-hydrogenase complex (Berney et al., 2014). In *E. coli*, homologous proteins are involved in the conversion of formate to CO_2_ and H_2_ under conditions of anaerobic respiration in the absence of an external terminal electron acceptor (Leonhartsberger et al., 2002). Facultative H_2_ metabolism is central for mycobacterial persistence. Mycobacteria enhance long-term survival by up-regulating hydrogenases during energy and oxygen limitations (Greening and Cook, 2014).

*Rv1620c-Rv1622c* (*cydC, D, B* respectively) encode proteins, which are involved in the cytochrome biogenesis and active transport across the membrane of components involved in the assembly of cytochrome. The expression of *CydDC* is linked to the incorporation of heme cofactors into a variety of periplasmic cytochromes, as well as the bd-type respiratory oxidases. CydB is the component of the aerobic respiratory chain that is supposedly predominant when cells are grown at low aeration, and is up-regulated under low pH (Baker et al., 2014). It has been reported that the presence of bd-type oxidases is correlated with bacterial virulence. For example, growth of mycobacteria at low oxygen tensions enhances both the expression of a bd-type oxidase and cell invasion (Bermudez et al., 1997).

Down-regulated genes belong to different functional categories. Among them, three genes with chaperone functions attract special attention. All these genes are essential for *M. tuberculosis* growth *in vitro* (Griffin et al., 2011; Sassetti et al., 2003). *Rv0440* encodes GroEL2, the chaperone belonging to the HSP60 family. Its chaperone-like functions provide resistance to stress (Qamra et al., 2004) and modulate host immune responses (Lewthwaite et al., 2007; Naffin-Olivos et al., 2014). GroEL2 is highly induced in response to environmental cues during infection like heat shock, oxidative stress, growth in macrophages and hypoxia (Qamra et al., 2005). The HupB protein encoded by *Rv2986c* belongs to the histone-like family of prokaryotic DNA-binding proteins capable of wrapping DNA to stabilize it, and prevent DNA denaturation under extreme environmental conditions (Kumar et al., 2010). It is involved in controlling the transfer of mycolic acids to sugars by the Ag85 complex (Katsube et al., 2007), as well as siderophore biosynthesis, and is essential for mycobacteria growth in macrophages (Pandey et al., 2014).

WhiB2 encoded by *Rv3260c* belongs to the WhiB family of transcriptional regulators. Its apo-form displays a chaperone activity, preventing aggregation and providing correct refolding of proteins; this activity does not require ATP and is independent of its own oxidized or reduced status and co-chaperones (Konar et al., 2012). The homologue of whiB2 in *M. smegmatis*, WhmD, participates in septa formation during cell division. WhmD overexpression decreases the linear size of *M. smegmatis* cells (Raghunand and Bishai, 2006). In addition, the newly formed cell walls are more susceptible to lysis (Gomez and Bishai, 2000). It was suggested that WhiB2 is involved in the assembly and stabilization of the FtsZ ring around the cell septum during division (Huang et al., 2013).

Two other down-regulated genes, *Rv2987c* and *Rv2988c*, encode subunits of putative 3-isopropylmalate dehydratase and are involved in leucine biosynthesis. *In vivo* and *in vitro* studies demonstrate that leucine-auxotrophic *M. tuberculosis* strains do not replicate inside host cells (Hondalus et al., 2000).

Overall, up-regulated genes fall into “intermediate metabolism and respiration” functional category, and either belong to the DosR regulon directly (*Rv0079-Rv0081*), or are connected through DosR-regulated transcriptional factor *Rv0081* (*Rv0082-Rv0087*). All up-regulated genes are thought to be involved into mycobacterial survival inside macrophages.

The list of down-regulated genes is functionally more diverse. Of interest is a decreased of genes encoding chaperone proteins, such as GroEL2, HupB, WhiB2. Theoretically, their expression should be rather increased under unfavorable conditions. To find an explanation of this paradox, we studied TB databases (http://genome.tbdb.org), concentrating on groups of genes co-expressed with these chaperones genes. It appeared that the majority of DEGs, including g*roEL2, hupB* and *whiB2*, are co-expressed with genes of the functional category J: Translation, ribosomal structure and biogenesis (Clusters of Orthologous Groups, COG, (Tatusov et al., 2000)). In all cases, irrespective to up- or down-regulation of DEGs, there was reversed correlation with genes of the J category (Table 2). These data suggest that the MTS1338 overexpression leads to transcriptional changes that correlate with a translation slowdown.

**Table 2.** DEGs, and their correlation with expression of different functional categories (according to COG) abbreviations: C – energy production and conversion; J – Translation, ribosomal structure and biogenesis; K – transcription; M – cell envelope biogenesis, outer membrane; O – posttranslational modifications, protein turnover, chaperones; T – signal transduction mechanisms. Orange boxes stand for up-regulated DEGs, green – down-regulated DEGs.

| gene |  | product | Genes with correlated expression, categories enriched |  |
| --- | --- | --- | --- | --- |
|  |  |  | Positive correlation | Negative correlation |
| *Rv0079* | *Rv0079* |  | **T** |  |
| *Rv0080* | *Rv0080* |  | **T** | **C, J** |
| *Rv0087* | *hycE* | Possible formate hydrogenase HycE |  | **J** |
| *Rv0440* | *groEL2* | 60 kDa chaperonin 2 GroEL2 | **J** |  |
| *Rv1158c* | *Rv1158c* | Conserved hypothetical ala-, pro-rich protein | **C, M** |  |
| *Rv1620c* | *cydC* | Probable 'component linked with the assembly of cytochrome' transport transmembrane ATP-binding protein ABC transporter CydC |  | **J** |
| *Rv1621c* | *cydD* | Probable 'component linked with the assembly of cytochrome' transport transmembrane ATP-binding protein ABC transporter CydD |  | **J** |
| *Rv1622c* | *cydB* | Probable integral membrane cytochrome D ubiquinol oxidase (subunit II) CydB (cytochrome BD-I oxidase subunit II) |  | **J** |
| *Rv2947c* | *pks15* | Probable polyketide synthase Pks15 | **J** | **K** |
| *Rv2986c* | *hupB* | DNA-binding protein HU homolog HupB (histone-like protein) | **J** |  |
| *Rv2988c* | *leuC* | Probable 3-isopropylmalate dehydratase (large subunit) LeuC (isopropylmalate isomerase) (alpha-IPM isomerase) (IPMI) | **K** |  |
| *Rv3019c* | *esxR* | Secreted ESAT-6 like protein EsxR (TB10.3) (ESAT-6 like protein 9) | **C, O** |  |
| *Rv3136* | *PPE51* | PPE family protein PPE51 | **J** |  |

Summarizing, our results suggest an important potential role of MTS1338 in pathogenesis of mycobacteria-triggered diseases. An increase in MTS1338 production during infection *in vivo* and in activated macrophages, changes in the expression of genes important for mycobacterial metabolism and a better survival under low pH accompanying MTS1338 overexpression – all suggest that this sRNA may well contribute to successful persistence of *M. tuberculosis* within host cells.

## Supporting information

Oligonucleotides used in the study

RNA-seq data

qPCR validation of RNA-seq

## Acknowledgements

This work was supported by the Russian Science Foundation grant №18-15-00332 to TA (new MTb strains, RNA-seq experiments and analyses, stresses in vitro); grant №18-45-04015 to AA (in vivo and ex vivo infection experiments)

## Authors contribution

Conceived the idea and designed the experiments: EGS, DI, AK, AA, TA. Performed the experiments: EGS, AG, YS, KM, OB, AO, NL. Analyzed the data: EGS, AK, AA, TA. Wrote the paper: EGS, AK, AA, TA.

## Data availability statement

The data sets supporting the results of this article are available in the GEO data repository under the accession number GSE137857.

## Supplementary material

**Supp Table 1. Oligonucleotides used in the study**

**Suppl Table 2. RNA-seq data**

