## Supplementary material for "MTS1338, a small *Mycobacterium tuberculosis* RNA, regulates transcriptional shifts consistent with bacterial adaptation for entering into dormancy and survival within host macrophages": Oligonucleotides used in the study

**Supplementary Table 1. Oligonucleotides used in the study**

| Name | Sequence |
| --- | --- |
| MTS1338-f | ACCGGGGAAACCCGGTGAT |
| MTS1338-r | AACAGGATGAGGATCTGCCC |
| MTS1338 HindIII-f | TATTCGAAGGGGAAACCCGGTGATCT |
| MTS1338 HindIII-r | TATTCGAAAACAGGATGAGGATCTGCCC |
| qPCR_MTS1338-f | GTGCTGGGCGATTGAGC |
| qPCR_MTS1338-r | GCGGTAGCCCCGTCTT |
| qPCR_16S-f | TACGTAGGGTGCGAGCGTTG |
| qPCR_16S-r | CCCGCACGCTCACAGTTAAG |
| qPCR_Rv0081-f | GCCTGGAGTCGTCGAACCT |
| qPCR_Rv0081-r | GGGTGCGGCAATCGAATAGAT |
| qPCR_Rv0083-f | CGTTTCTGCTGGCGTGGGA |
| qPCR_Rv0083-r | CAACACCACCAGCCCGAC |
| qPCR_Rv0086-f | TTTGGGTAGCGATCGAGGCCA |
| qPCR_Rv0086-r | TACCCAAGAAGGCGACGGC |
| qPCR_Rv1621-f | GCTGGCTACCACTAACCCCTC |
| qPCR_Rv1621-r | TTCCGCGATGCGTTGTTCC |
| qPCR_Rv2986-f | CGTCACCATTACCGGGTTCG |
| qPCR_Rv2986-r | CACAACCGCTTTGAATTGCGC |
| qPCR_Rv3136-f | AAACAGCGATCCAAGCCAGG |
| qPCR_Rv3136-r | CCGCAGTGTTCTGGCCGA |
