## Supplementary material for "MTS1338, a small *Mycobacterium tuberculosis* RNA, regulates transcriptional shifts consistent with bacterial adaptation for entering into dormancy and survival within host macrophages": qPCR validation of RNA-seq

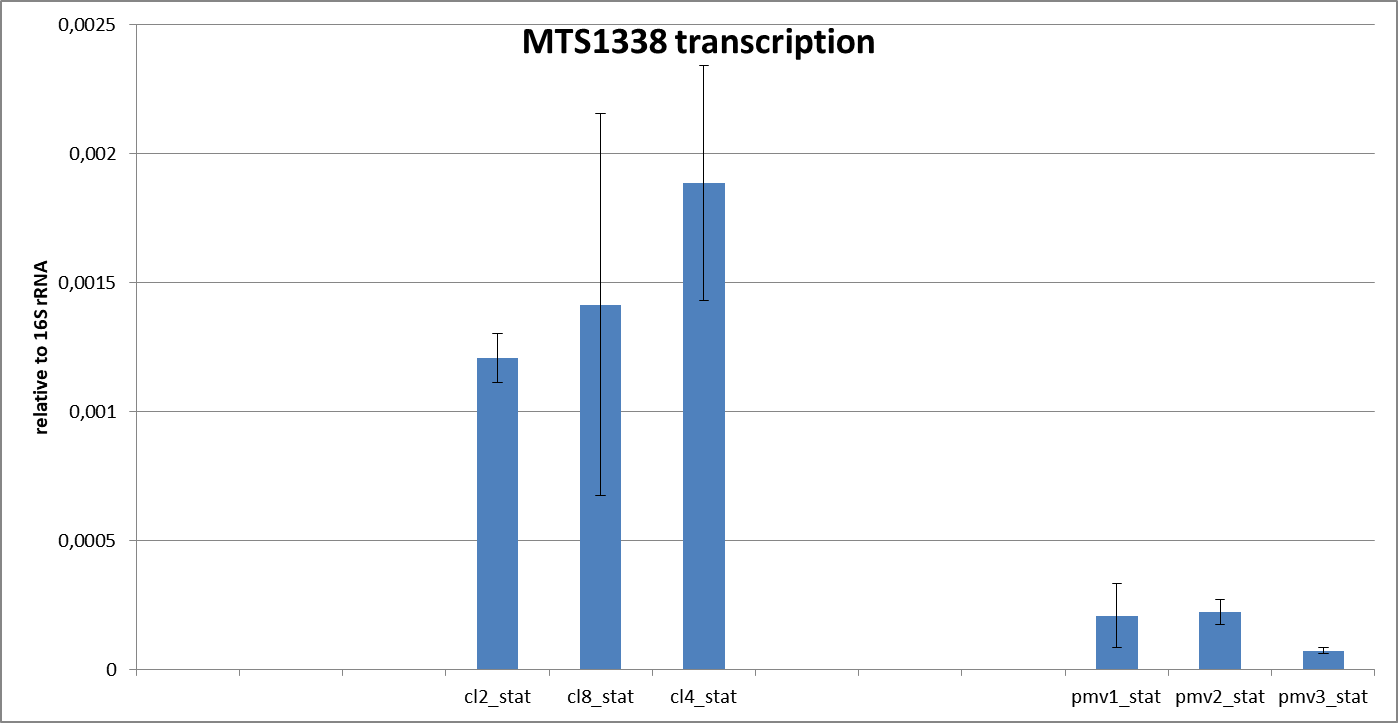

Figure 1A. MTS1338 transcription level in samples used for RNA-seq. pMV stands for control *M.tuberculosis* strains. Cl2, cl8, cl4 – OVER strains, independent clones.

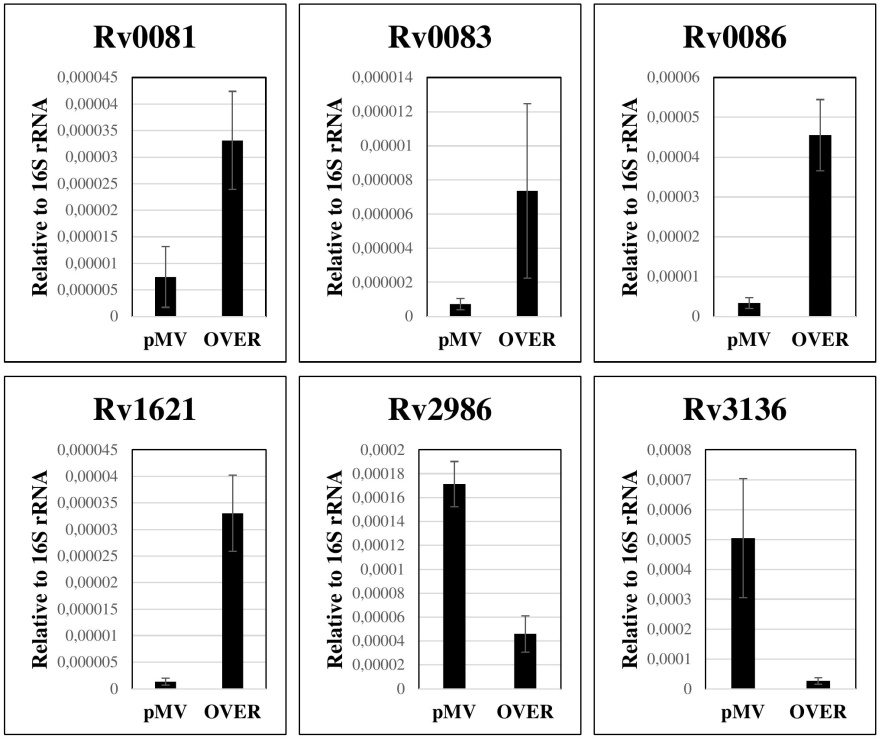

Figure 1B. Validation of RNA-seq data by qRT-PCR
